# Bioherbicidal activity and host range of *Teratoramularia rumicicola* strain TR4 isolated from *Rumex crispus* in Japan

**DOI:** 10.1101/2025.08.06.668795

**Authors:** Masataka Izumi, Toyozo Sato

## Abstract

*Rumex* species, including *R. crispus*, *R. obtusifolius*, and *R. japonicus*, are problematic weeds that reduce forage production and hinder effective land use in Japanese pasture systems. This study aimed to identify a bioherbicide candidate that selectively targets *Rumex* weeds while remaining safe for forage crops. To this end, we isolated a fungal pathogen from a naturally infected *R. crispus* leaf collected in Japan and conducted morphological and molecular identification and host range evaluations. The isolate, designated TR4, was identified as *Teratoramularia rumicicola* based on its morphological characteristics and phylogenetic analysis of concatenated internal transcribed spacer (ITS) and large subunit (LSU) rDNA sequences. Host range studies revealed that TR4 caused visible symptoms and significantly reduced shoot biomass in all three *Rumex* species, while no disease symptoms were observed in any of the five forage crops tested. This study reports *T. rumicicola* for the first time in Japan, extending its known distribution beyond South Korea. The host specificity and liquid culture compatibility of TR4 suggest its potential as a locally adapted microbial herbicide for selective *Rumex* control in pasture systems.

## Introduction

Herbicides have long been used to enable efficient weed control in modern agriculture. However, the repeated use of herbicides has led to the widespread emergence of herbicide-resistant weed biotypes (Heap 2025). Moreover, the discovery of herbicides with novel modes of action has slowed significantly in recent decades (Umetsu and Shirai 2020), increasing the urgency to develop alternative weed management strategies. Integrated Weed Management, which incorporates chemical, mechanical, cultural, and biological approaches, is gaining prominence as a sustainable approach to weed control (Westwood et al. 2018), as it reduces the selection pressure associated with sole dependence on chemical herbicides.

Among the various components of Integrated Weed Management, microbial herbicides, which are applied in large quantities to achieve rapid weed suppression via an inundative approach, have gained renewed attention as alternatives to synthetic herbicides (Duke 2024). They offer distinct advantages, including high crop safety, low toxicity to mammals and nontarget organisms, limited environmental persistence, relatively low development and registration costs, and no reported cases of resistance (Duke et al. 2022). However, their broader adoption has been constrained by challenges such as inconsistent efficacy under diverse field conditions, the need for low-cost mass production, development of formulations that enhance shelf life and field performance, and assurance of biosafety throughout the product lifecycle (Gressel 2024).

*Rumex* species are potential targets for the development of microbial herbicides, particularly in temperate forage systems. *R. obtusifolius* L. and *R. crispus* L. are among the most problematic broadleaf weeds in temperate pasturelands and arable fields, owing to their perennial growth and ability to compete with forage crops and reduce productivity (Zaller 2004). Notably, *R. obtusifolius* has evolved resistance to acetolactate synthase-inhibiting herbicides, such as florasulam, metsulfuron-methyl, and thifensulfuron-methyl, in France (Heap 2025). In parallel with chemical control efforts, several fungal pathogens, including *Uromyces rumicis*, *Ramularia rubella*, *Venturia rumicis*, and *Armillaria mellea*, have been investigated as biological control agents for *Rumex* species, although their practical use remains limited owing to insufficient efficacy and biosafety concerns (Bond et al. 2007). These limitations highlight the need to identify novel microbial candidates with greater potential for practical applications.

In this study, we focused on Japan, where *R. obtusifolius*, *R. crispus*, and *R. japonicus* Houtt. are widely distributed in pastures and disturbed habitats. While the former two are introduced species and *R. japonicus* is native, all three are recognized as invasive weeds and are listed as “alien species of concern” by the Ministry of the Environment (Ministry of the Environment, Japan 2015). In addition, *R. obtusifolius* and *R. japonicus* are designated as “prohibited weed species” by the Japan Grassland Agriculture and Forage Seed Association because of their significant impact on forage production, and their contamination is not tolerated in seed certification (Japan Grassland Agriculture and Forage Seed Association 2019). These facts underscore the serious weed problems posed by *Rumex* species in Japan; however, the available chemical herbicides for their control remain limited, such as thifensulfuron-methyl. Considering the global development of herbicide resistance in *Rumex* species, alternative control measures, such as microbial herbicides, are urgently needed. However, to our knowledge, no studies have been published on microbial herbicides targeting *Rumex* weeds in Japan.

To address this gap, we aimed to isolate and identify a fungal pathogen with bioherbicidal activity against *Rumex* species of relevance to forage systems in Japan. A candidate strain was obtained from a naturally infected *R. crispus* leaf. Morphological and molecular analyses were conducted to determine its taxonomic identity, and host range tests were performed to evaluate its potential as a selective microbial herbicide.

## Materials and Methods

### Fungal Isolation

A symptomatic leaf of *Rumex crispus* was collected from a roadside plant in Tsukuba City, Ibaraki Prefecture, Japan on 17^th^ November 2024. The leaf was detached and incubated at 25 °C under moist and light conditions for 12 days until conidial structures emerged from the lesions. Emerging conidia were collected and streaked onto potato dextrose agar (PDA, BD Difco^TM^/BBL^TM^) amended with lactic acid for five days for single-conidium isolation. The plates were incubated at 25 °C for fungal growth. One isolate, designated as TR4, was selected as the representative isolate for subsequent experiments. The isolate TR4 was deposited as MAFF 248116 in the NARO Genebank (National Agriculture and Food Research Organization, Japan).

### Morphological Observations

TR4 was transferred to the center of PDA plates (90 mm in diameter) with a small piece of mycelium and incubated at 25 °C in the dark. After one month, colony characteristics, including growth rate, surface texture, pigmentation on both the surface and reverse sides, and margin shape were recorded. Conidial suspensions were prepared by flooding the colony surface with sterile distilled water, and mounted on sterile distilled water. Conidial length and septation were observed using a light microscope (BH-2, Olympus, Japan). Morphological characteristics were compared with reference to the published descriptions of *Teratoramularia rumicicola* S.I.R. Videira, H.D. Shin & P.W. Crous and *Teratoramularia rumicis* Kushwaha, S.K. Verma, S. Yadav & Raghv. Singh, known foliar pathogens of *Rumex* species (Videira et al. 2016; Verma et al. 2021).

### DNA Extraction

TR4 was cultured on PDA plates at 25°C in the dark for six weeks. The mycelia were harvested and immediately frozen in liquid nitrogen. The frozen material was pulverized using a bead homogenizer (MB456KU(S), Yasui Kikai, Japan) at 2000 rpm for 30 seconds. Genomic DNA was extracted using the NucleoSpin Soil Kit (Macherey-Nagel, Germany) according to the manufacturer’s protocol. DNA quality and concentration were assessed using a NanoDrop spectrophotometer (NanoDrop 2000, Thermo Fisher Scientific, USA).

### PCR Amplification and Sequencing

PCR amplification was performed for two genomic regions: the internal transcribed spacer (ITS) region using primers ITS1 (5’-TCCGTAGGTGAACCTGCGG-3’) and ITS4 (5’-TCCTCCGCTTATTGATATGC-3’) (White et al. 1990), and the large subunit ribosomal RNA (LSU) region using primers LR0R (5 ’-ACCCGCTGAACTTAAGC3’) and LR7 (5’-TACTACCACCAAGATCT-3’) (Vilgalys and Hester 1990). These primer sets were selected with reference to Verma et al. (2021) and Choi et al. (2025), who employed ITS and LSU sequences to discriminate *T. rumicicola* and *T. rumicis*.

Each 10 µL PCR reaction contained 5.0 µL of 2× PCR buffer, 2.0 µL of 2 mM dNTPs, 0.3 µL of each primer (10 µM), 0.2 µL of KOD FX Neo DNA polymerase (Toyobo, Japan), 1.2 µL of sterile distilled water, and 1 µL of the genomic DNA. The thermal cycling conditions were as follows: initial denaturation at 94°C for 2 minutes; 35 cycles of 98°C for 10 seconds, 58°C for 15 seconds, and 68°C for 30 seconds; and a final extension at 68°C for 5 minutes. Successful amplification was confirmed by electrophoresis on a 1.5% agarose gel stained with SYBR Green (Takara Bio, Japan).

The PCR products were purified using ExoSAP-IT (Thermo Fisher Scientific, USA) and submitted to Eurofins Genomics (Tokyo, Japan) for Sanger sequencing. Chromatograms were manually assembled and edited using CodonCode Aligner version 12.0.1. The resulting consensus sequences were analyzed using the BLAST n algorithm on the NCBI website to determine their closest taxonomic affiliation. The sequences were deposited in GenBank under accessions PX069134 (ITS) and PX069135 (LSU).

### Phylogenetic Analysis

ITS and LSU sequences obtained in this study were concatenated and included in the phylogenetic analysis. Reference sequences of *T. rumicicola*, *T. rumicis*, *T. infinita*, *T. kirschneriana*, and *T. persicariae* were retrieved from GenBank. The reference sequences and their GenBank accessions are listed in Supplementary Table 1. Multiple sequence alignment was conducted using MUSCLE in MEGA version 11 (Tamura, Stecher, and Kumar 2021), and ambiguously aligned and highly variable terminal regions were manually trimmed. The final concatenated alignment of ITS and LSU sequences consisted of 1,191 bp.

Maximum likelihood analysis was performed using RAxML v8.2.4 under the GTR+GAMMA model with 1000 rapid bootstrap replicates (Stamatakis 2014). The tree was rooted with *Staninwardia suttonii* CPC 13055 and visualized using Interactive Tree of Life (Letunic and Bork 2024). Bootstrap values ≥70% were displayed.

### Plant Materials and Growth Conditions

Seeds of the weed species were obtained by purchase (*R. crispus* and *R. obtusifolius*, Espec Mic Co., Ltd., Japan) or by field collection in Kyoto City, Kyoto Prefecture (*R. japonicus*). Seeds of the forage species were obtained from commercial suppliers: *Dactylis glomerata* and *Lolium multiflorum* from Snow Brand Seed Co., Ltd. (Japan), *Phleum pratense* and *Trifolium repens* from Takii & Co., Ltd. (Japan), and *Medicago sativa* from Kaneko Seeds Co., Ltd. (Japan).

All seeds were pre-germinated on moistened filter paper in Petri dishes at 25°C/15°C under a 12 h light/12 h dark photoperiod in a growth chamber. After several days, the germinated seedlings were transplanted into plastic pots (φ = 5 cm) filled with a 1:1 (v/v) mixture of vermiculite and akadama soil (Japanese granular clay soil) and grown under the same conditions. One seedling was grown per pot and watered as needed. Inoculations were performed at the 1–2 leaf stage, approximately 2 weeks after sowing for forage species and 3 weeks for weed species.

### Inoculum Preparation and Application Conditions

For preculture, a 6-mm-diameter agar disc of isolate TR4 grown on PDA was excised and inoculated into 50 mL of 0.5% YM liquid medium, consisting of 0.5% yeast extract and 0.5% malt extract (both from Solabia Biokar Diagnostics, France), and 0.5% D-(+)-glucose (Nacalai Tesque, Japan), in a 200 mL Erlenmeyer flask. The culture was incubated at 25 °C for 3 -4 days with rotary shaking at 130 rpm using a rotary shaker (NR-20, TAITEC Corporation, Japan). Subsequently, 1.5 mL of the preculture was transferred into 150 mL of 0.5% YM liquid medium in a 500 mL Sakaguchi flask. The culture was incubated at 25 °C for 7 days with reciprocal shaking at 130 rpm in a shaking water bath incubator (BT200, Yamato Scientific Co., Ltd., Japan).

The culture broth was filtered through two layers of cotton gauze (High Gauze White, Iwatsuki Co., Ltd., Japan) to remove the large mycelial fragments. The resulting suspension was adjusted with YM liquid medium to a final concentration of 3.0 × 10^5^ propagules/mL for weed species and 1.0 × 10^6^ propagules/mL for forage species using a hemocytometer. The term “propagules” refers to both conidia and hyphal fragments, although their proportions were not determined in this study. The suspension was applied to *R. crispus*, *R. japonicus*, *R. obtusifolius*, *D. glomerata*, *L. multiflorum*, *P. pratense*, *M. sativa*, and *T. repens* at the 1-2 leaf stage using a handheld sprayer (Daiya Spray, Furupla Co., Ltd., Japan) at an application volume equivalent to 4000 L/ha. YM liquid medium was used as a non-treated control. Following treatment, plants were enclosed in moistened polyethylene bags at 25°C in the dark for 48 hours. They were transferred to growth conditions at 25 °C under a 12 h light/12 h dark photoperiod, while remaining enclosed in moistened polyethylene bags.

### Biological Evaluation

Leaf-level disease incidence was assessed in all test species 14 days after treatment (DAT) and again at 21 DAT for *Rumex* species. Leaf-level disease incidence was calculated as the number of leaves exhibiting visible symptoms divided by the total number of leaves. Leaves were considered symptomatic when white conidiomata, characteristic of *T. rumicicola* (Choi et al. 2025), was observed on the surface. Cotyledons that withered naturally without visible fungal colonization, which were observed in control plants, were excluded from both the symptom count and total leaf number to ensure that only pathogen-induced symptoms were recorded. Surface symptoms on infected *Rumex* leaves were documented using a digital microscope (VHX-970F, KEYENCE, Japan). In addition, the aboveground parts of *Rumex* species were harvested at 21 DAT and immediately weighed to determine shoot fresh weight. To confirm pathogen re-isolation as part of Koch’s postulates, conidia were collected from symptomatic *Rumex* leaves at 21 DAT using a sterile platinum loop and reinoculated onto PDA plates. After several days of incubation at 25 °C, colony and conidial morphologies were examined and compared with those of the original isolate.

### Statistical Analysis

All bioassay results are presented as the mean ± standard error from at least three biological replicates. Statistical analyses were performed using the Mann–Whitney U test for leaf-level disease incidence, and Student’s *t*-test for shoot biomass after confirming homogeneity of variances by an F-test in R (version 4.5.0). Each bioassay was independently repeated at least twice to ensure its reproducibility.

## Results and Discussion

### Isolation of the Fungal Pathogen

A symptomatic leaf of *Rumex crispus* was collected from a roadside plant in Tsukuba, Japan. After incubation under moist conditions at 25 °C for 12 days, lesions with dark purple to brown margins and white fungal growth were observed on the leaf surface (Figure 1a, b). Microscopic observation revealed the presence of twiggy conidiophores and cylindrical conidia (Figure 1c). Conidia were transferred to lactic acid-amended PDA for isolation, and one isolate, designated as TR4, was selected for further study.

**Figure 1.**
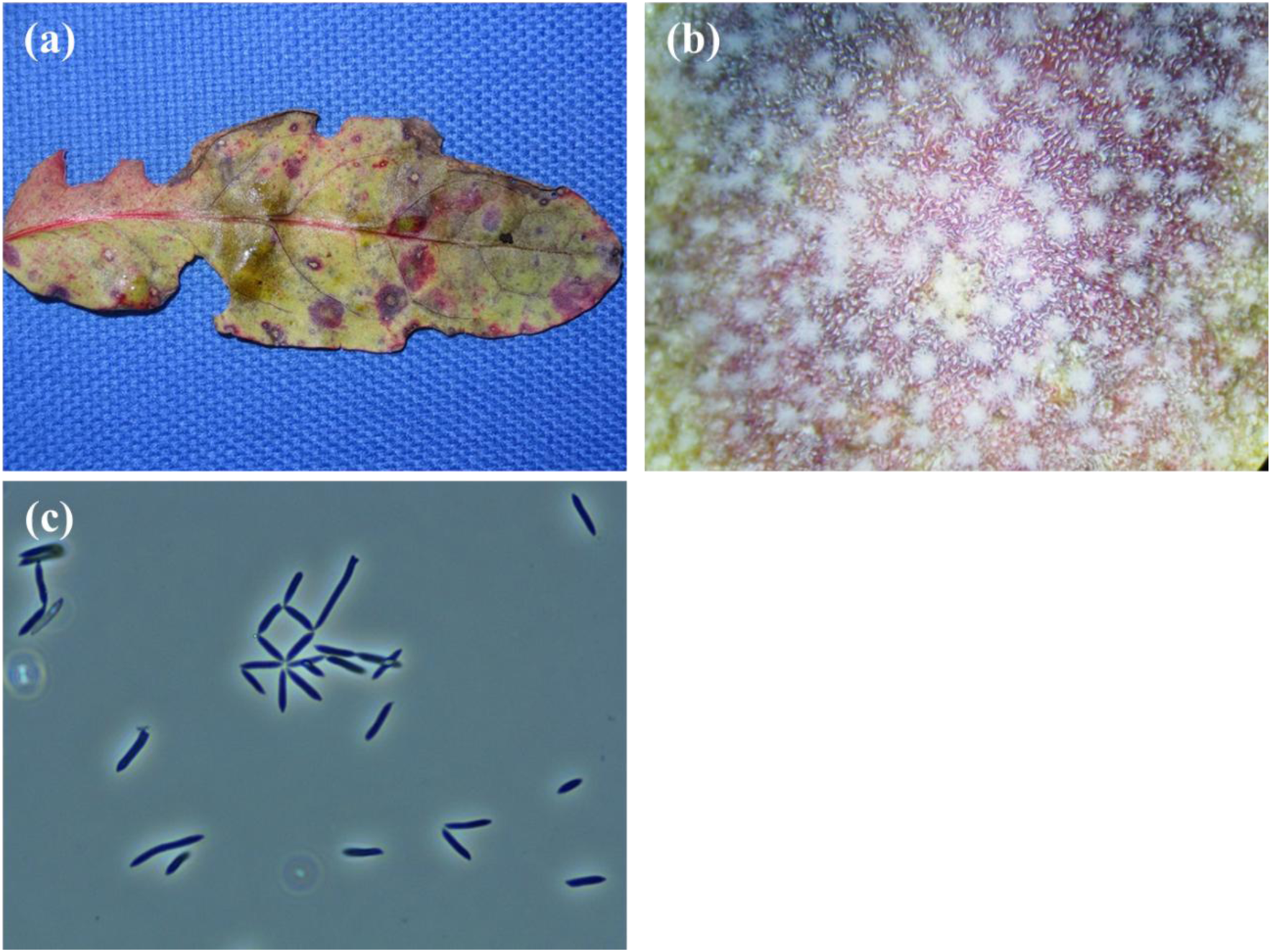
Isolation of TR4 from a naturally infected leaf of *Rumex crispus*. (a) Symptomatic leaf after 12 days of incubation at 25°C under moist and light conditions. (b) Lesion surface observed under a stereomicroscope, with visible white conidiophores and conidia. (c) Light microscopy image of conidia collected from the lesion (×400), showing cylindrical morphology (phase contrast microscopy).

### Morphological Characterization

Colonies on PDA incubated at 25 °C in the dark grew very slowly, reaching approximately 10 mm in diameter after one month (Figure 2). They were subcircular with moderately elevated, uneven surfaces and coarse, wrinkled textures. The surface appeared dark, with white zones at the center and margin. The surrounding medium was faintly stained red due to pigment diffusion, particularly toward the colony center. On the reverse, colonies showed a dark center and pale edge. Microscopic observation revealed hyaline, cylindrical conidia, mostly aseptate with occasional 1 -septate forms, measuring 5.5–21.0 × 1.7–2.9 μm (*n* = 30), with slightly rounded ends (Figure 3). These features closely resembled those of the ex-type strain CPC 14653 of *Teratoramularia rumicicola* (Videira et al. 2016). According to Verma et al. (2021), *T. rumicis*, the species most closely related to *T. rumicicola*, produces longer conidia with a higher frequency of septation (0 –2 septa) than *T. rumicicola*. This morphological distinction supports the identification of TR4 as *T. rumicicola*.

**Figure 2.**
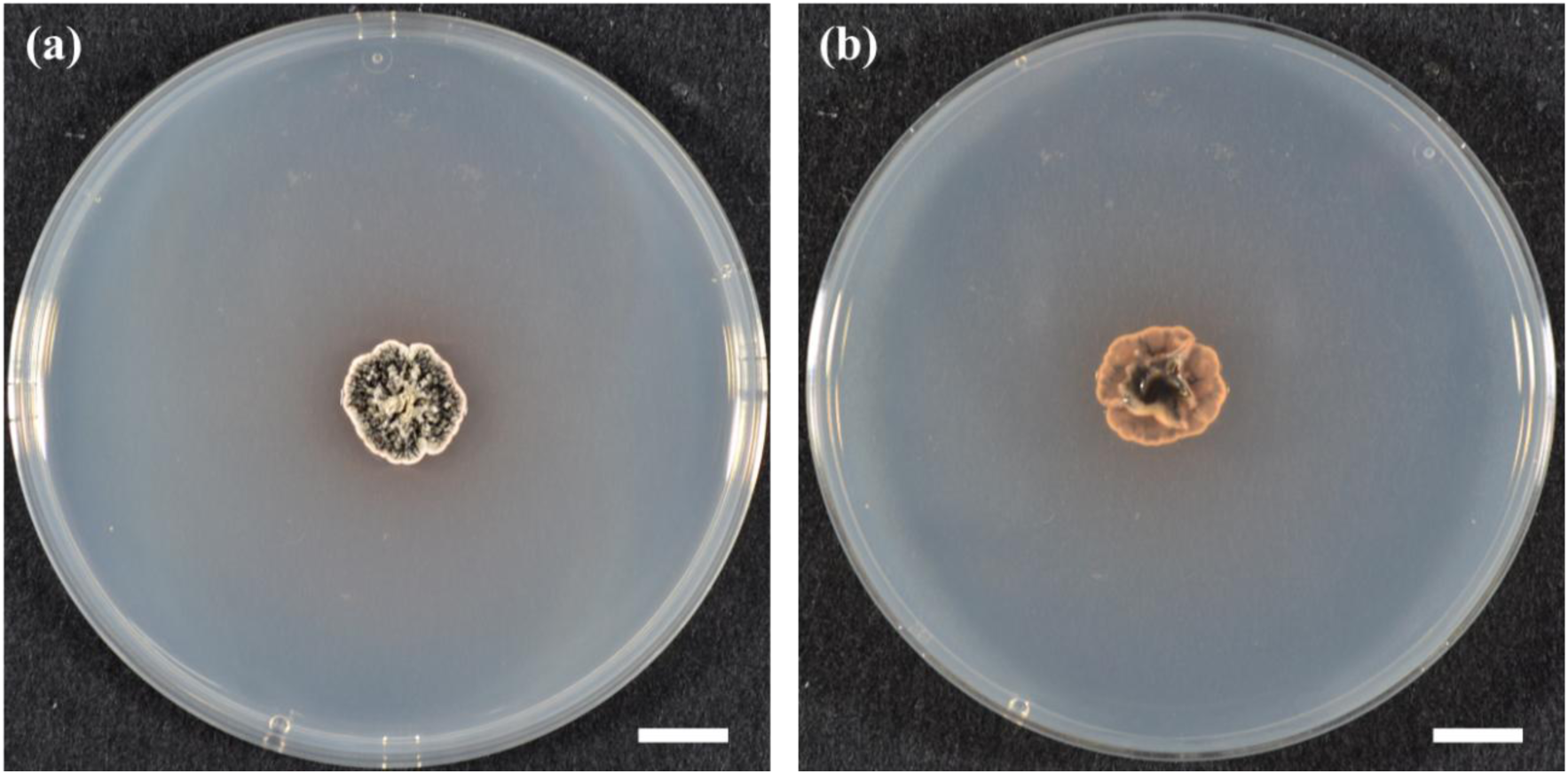
Colony Morphology of Isolate TR4 on PDA. Colony appearance of TR4 grown on potato dextrose agar (PDA) after one month of incubation at 25 °C in the dark: (a) surface view; (b) reverse view. Scale bars = 10 mm

**Figure 3.**
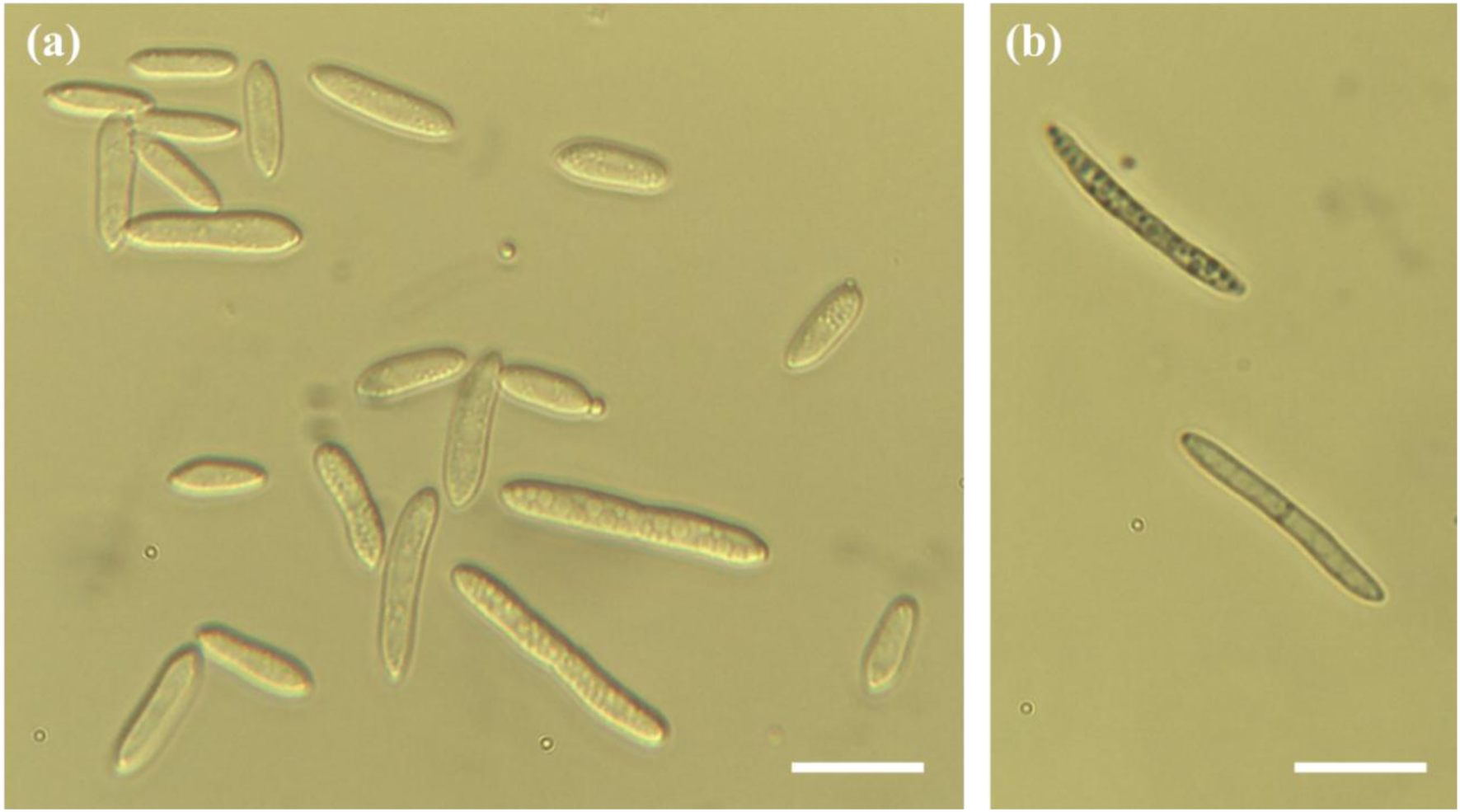
Microscopic Morphology of TR4 Conidia. Microscopic morphology of TR4 conidia observed under a light microscope. The conidia were mostly aseptate with occasional 1-septate forms, cylindrical, and hyaline. Scale bars = 10 µm

### BLAST and Phylogenetic Analysis

BLASTn analysis of the ITS sequence of TR4 showed 100% identity to three *T. rumicicola* accessions (OL711641, PP301328, and PP301329), as well as to the ex-type strain of *T. rumicis* (NR_174642). The ex-type strain of *T. rumicicola* (NR_154517) also showed a very high similarity, with 99.58% identity (469/471 bp). These results indicate that the ITS region alone lacks sufficient resolution to distinguish *T. rumicicola* from *T. rumicis*, highlighting the importance of additional markers for accurate species identification.

The LSU sequence of TR4 showed 98.6% identity (697/707 bp) to the ex-type strain of *T. rumicis* (MW276098), whereas 100% identity to the ex-type strain and two other *T. rumicicola* accessions (KX287254, KX287255, and KX287256). Two additional *T. rumicicola* accessions (PP301336 and PP301337) exhibited 99.86% identity (722/723 bp). These results demonstrate that LSU provides a higher resolution than ITS for distinguishing *T. rumicicola* from *T. rumicis* and supports the assignment of TR4 to *T. rumicicola*.

A phylogenetic tree was constructed based on concatenated ITS and LSU sequences using the maximum-likelihood method. The resulting tree resolved four well-supported clades corresponding to *T. kirschneriana*, *T. infinita*, *T. persicariae*, and a combined clade containing both *T. rumicis* and *T. rumicicola*. TR4 clustered in the same clade as the ex-type strains of both *T. rumicicola* and *T. rumicis*, along with other *T. rumicicola* reference strains, with a bootstrap support of 100%. Although both type strains were included within the same clade, the *T. rumicis* ex-type strain was positioned on a distinctly long branch, suggesting substantial divergence from TR4 and the *T. rumicicola* group. Previous studies by Verma et al. (2021) and Choi et al. (2025) reported *T. rumicicola* and *T. rumicis* as forming separate clades based on ITS–LSU concatenated analyses. The observed differences between our results and those of previous studies may, in part, be due to methodological choices, including alignment, trimming, or tree-building protocols. Nonetheless, the close phylogenetic proximity to the *T. rumicicola* group, distinct from the more divergent *T. rumicis* branch, is consistent with the identification of TR4 as *T. rumicicola*.

### Taxonomic Assignment of TR4

The colony characteristics and conidial morphology of TR4 closely resembled those of the *T. rumicicola* ex-type strain (CPC 14653). Molecular analyses further reinforced this identification, as TR4 exhibited complete identity in the LSU region and high similarity in the ITS region to the *T. rumicicola* ex-type strain, with phylogenetic placement among *T. rumicicola* reference strains. In addition to morphological and sequence-based evidence, the known geographic distribution supports this assignment. *T. rumicicola* has been isolated from multiple sites in South Korea, whereas *T. rumicis* is represented by a single isolate from India (Videira et al. 2016; Verma et al. 2021; Choi et al. 2025). Therefore, the Japanese origin of TR4 aligns more closely with that of *T. rumicicola*. Currently, only ITS and LSU sequences are available in GenBank for *T. rumicis*, and no additional gene regions (e.g., RPB2 and TEF1) have been reported. This lack of multi -locus data limits the potential for more robust species delimitations. Nonetheless, the integration of morphological, molecular, phylogenetic, and biogeographical evidence strongly supports the identification of TR4 as *T. rumicicola*.

### Host Range and Specificity

The leaf-level disease incidence at 14 DAT was 85.3 ± 6.5% in *R. obtusifolius*, 74.0 ± 7.3% in *R. crispus*, and 70.0 ± 4.2% in *R. japonicus*, all significantly higher than that in the control (*p* < 0.01, Mann–Whitney U test; Table 1). These values remained consistently high at 21 DAT. Infected leaves exhibited foliar necrosis and abundant white fungal growth, which are characteristic of *T. rumicicola* infections (Figure 5, S1, and S2). Shoot biomass at 21 DAT was significantly reduced in all *Rumex* species tested. The biomass of *R. crispus* and *R. japonicus* declined to 40.7 ± 6.5% and 45.2 ± 6.7% of the control, respectively, whereas *R. obtusifolius* retained a relatively higher biomass at 60.8 ± 7.8% (*p* < 0.01, Student’s *t*-test; Table 1). These results indicate a moderate but consistent suppression of shoot growth across species, with a slightly lower susceptibility in *R. obtusifolius*. Furthermore, conidia collected from symptomatic *Rumex* leaves inoculated with TR4 produced colonies and conidia that were morphologically identical to the original isolate. These observations fulfill Koch’s postulates and confirm the causal role of TR4 in symptom development.

**Figure 4.**
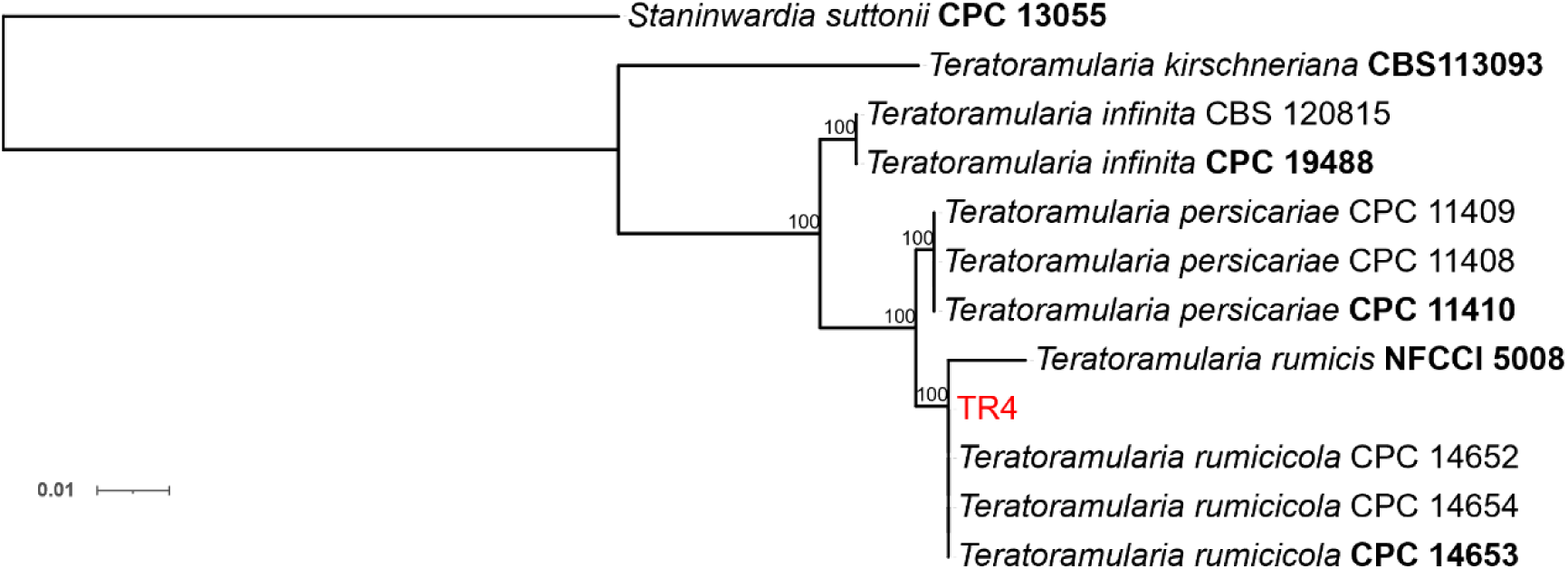
Phylogenetic Placement of Isolate TR4 Based on Concatenated ITS and LSU Sequences. Maximum likelihood tree constructed from concatenated ITS and LSU sequences using the GTR+GAMMA model. Bootstrap values ≥ 70% (1000 replicates) are shown at the nodes. TR4 is indicated in red font, and ex-type strains are indicated in bold font. Related *Teratoramularia* species were retrieved from GenBank. *Staninwardia suttonii* was used as the outgroup.

**Figure 5.**
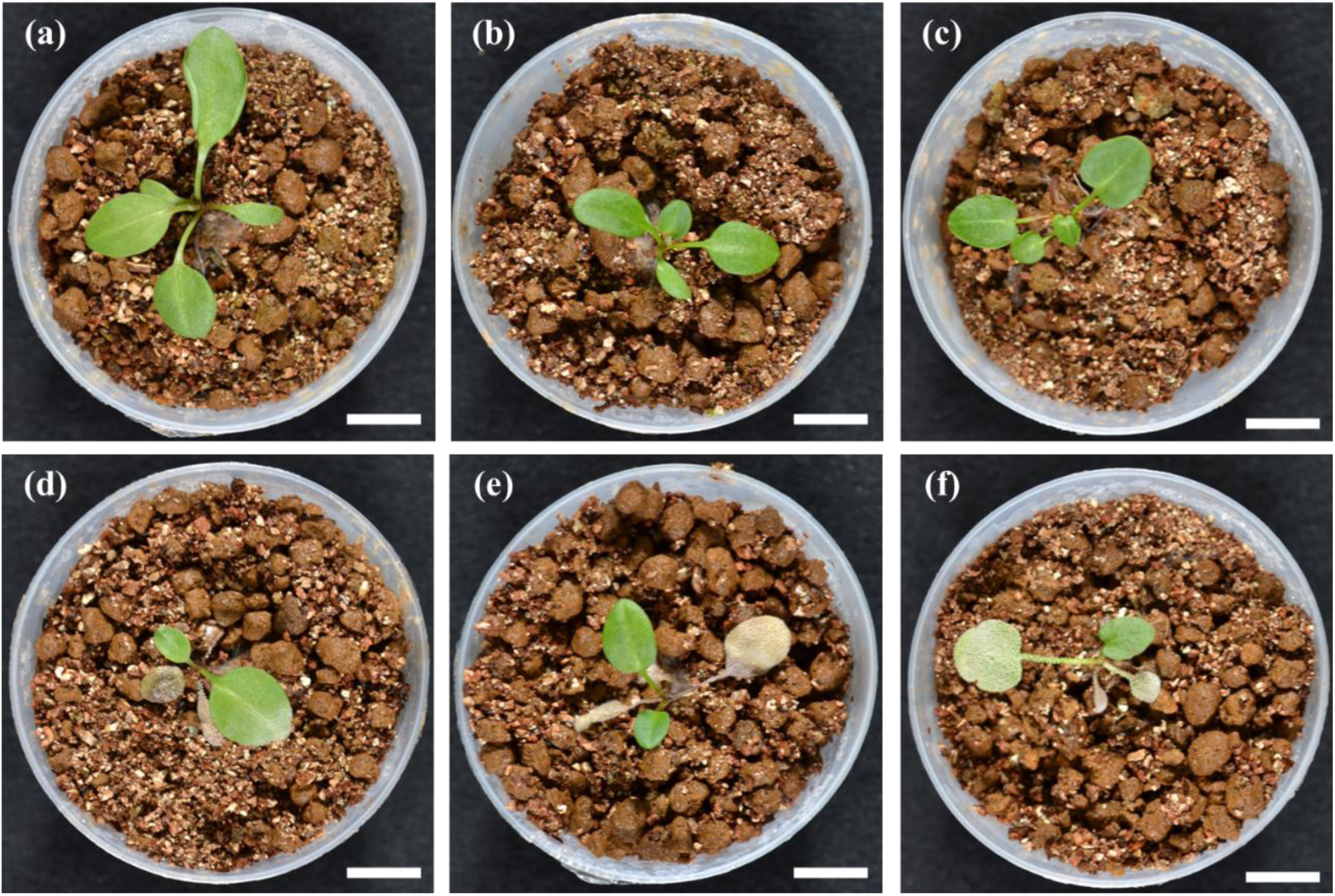
Pathogenicity of TR4 Against *Rumex* Species. Foliar appearance 21 days after inoculation. (a–c) Untreated control plants of *R. crispus*, *R. japonicus*, and *R. obtusifolius*, respectively. (d–f) Corresponding TR4-inoculated plants, showing visible white fungal growth and necrotic lesions. Scale bars = 10 mm

**Table 1.** Leaf-level Disease Incidence and Shoot Biomass of *Rumex* Weeds Treated with TR4. Values represent the mean ± SE (*n* = 4–5). Statistical differences from the control were determined using the Mann–Whitney U test or Student’s *t*-test (*p* < 0.01).

| Weeds | Leaf-level Disease Incidence (%) |  | Shoot Biomass (% of control) |
| --- | --- | --- | --- |
|  | 14 DAT | 21 DAT | 21 DAT |
| *Rumex crispus* | 74.0 ± 7.3 <sup>††</sup> | 72.0 ± 4.9 <sup>††</sup> | 40.7 ± 6.5 ** |
| *Rumex japonicus* | 70.0 ± 4.2 <sup>††</sup> | 69.7 ± 3.5 <sup>††</sup> | 45.2 ± 6.7 ** |
| *Rumex obtusifolius* | 85.3 ± 6.5 <sup>††</sup> | 83.0 ± 4.4 <sup>††</sup> | 60.8 ± 7.8 ** |

<sup>††</sup> $p < 0.01$ vs. control; Mann–Whitney U test, $n = 4-5$
\*\* $p < 0.01$ vs. control; Student's $t$ -test, $n = 4-5$

In contrast, the leaf-level disease incidence at 14 DAT was 0% in all five tested forage species. The primary diagnostic symptom was not observed on any of the leaf surfaces (Figure 6). These findings demonstrate that TR4 exhibits host specificity against *Rumex* weeds without affecting forage species, highlighting its potential as a selective bioherbicide in pasture systems.

**Figure 6.**
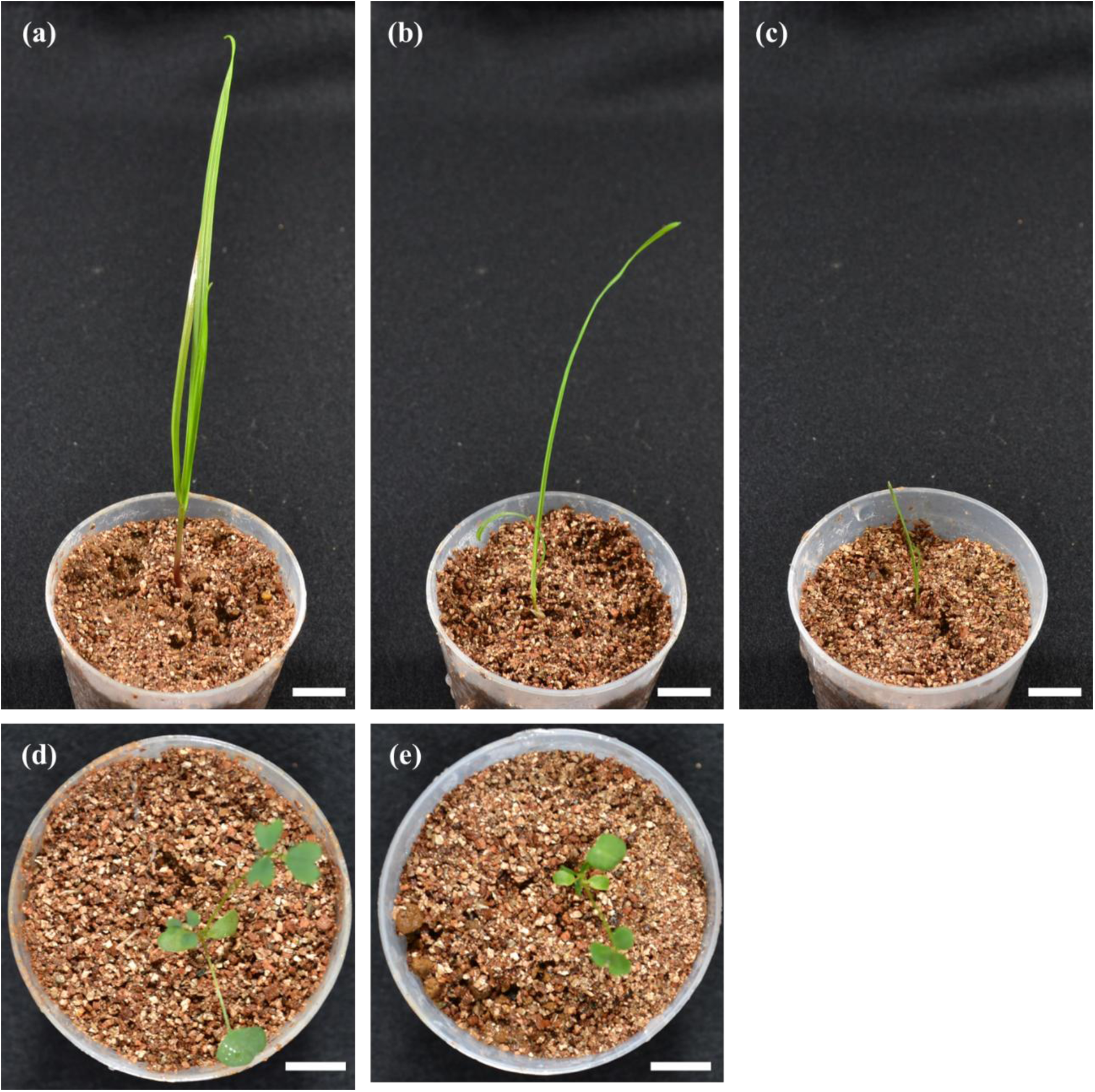
Lack of Pathogenicity of TR4 against Forage Species. Foliar appearance 14 days after inoculation. (a) *Lolium multiflorum*, (b) *Dactylis glomerata*, (c) *Phleum pratense*, (d) *Medicago sativa*, and (e) *Trifolium repens* following inoculation with TR4. No symptoms were observed in any of the species. Scale bars = 10 mm

### Significance and Implications of the Present Study

This study documents the first occurrence of *T. rumicicola* in Japan, thus extending its known geographic range beyond South Korea. Since its establishment in 2016, the genus *Teratoramularia* has remained poorly studied and *T. rumicicola* has been reported in only two publications, both based on South Korean isolates (Videira et al. 2016; Choi et al. 2025). This newly reported Japanese isolate provides valuable taxonomic and biogeographical insights. The consistency between morphological and molecular features further strengthens the framework for species identification within this understudied genus.

In addition to its original host *R. crispus*, TR4 exhibited pathogenicity toward the invasive alien species *R. obtusifolius* and the native weed *R. japonicus*, all of which are considered major weeds in Japanese pasture systems. In contrast, no pathogenic effects were observed in the five tested forage species, including grasses and legumes. These results from the preliminary screening provide the first experimental evidence that *T. rumicicola* strain TR4 may possess a host range and specificity desirable for bioherbicidal use. However, further studies are needed to evaluate its weed control efficacy under field conditions and confirm its safety toward a broader range of forage and crop species.

Although not directly examined in this study, TR4’s ability to grow in liquid culture suggests the potential for large-scale production. In contrast to obligate pathogens such as *Uromyces rumicis*, TR4 may offer a more practical platform for product development in terms of commercialization. Moreover, as a strain isolated locally in Japan, TR4 may exhibit greater ecological compatibility with the domestic environment and could be more amenable to regulatory approval as a regionally adapted microbial control agent. Further investigation is needed to optimize mass - production systems and assess the natural distribution and ecological behavior of *T. rumicicola* in Japan.

Collectively, this study contributes to the taxonomic understanding of the little-studied genus *Teratoramularia* and the early stage evaluation of *T. rumicicola* strain TR4 as a candidate microbial herbicide. By documenting its pathogenicity and host specificity, particularly in relation to *Rumex* weeds and non-target forage species, this study establishes a foundation for future development and deployment in weed management in pasture systems.

## Conclusion

This study reports the first isolation of *Teratoramularia rumicicola* in Japan and demonstrates its pathogenicity toward *Rumex crispus*, *R. obtusifolius*, and *R. japonicus*. No pathogenic effects were observed in the tested forage crops. TR4’s ability to grow in liquid culture suggests its potential for scalable production, laying the groundwork for future development as a regionally adapted microbial herbicide.

## Acknowledgements

This work was supported in part by the Research Initiative Grants Programme No. 2501 from the Weed Science Society of Japan (WSSJ), awarded to M. I. Language refinement and structural editing were supported by ChatGPT (OpenAI, GPT-4o) and Paperpal (provided by Editage). All AI-generated content was critically reviewed and revised by the authors.

## Author Contributions

M.I. conceived and led the study, performed most of the experiments, and wrote the manuscript. T.S. contributed to fungal isolation and provided expertise in inoculum preparation and fungal identification. All authors contributed to the critical revision of the manuscript, approved the final version to be published, and agreed to be accountable for all aspects of the work.

## Disclosure Statement

The authors declare no conflicts of interest.

## Data Availability Statement

The data that support the findings of this study are available from the corresponding author upon reasonable request.

## Figure Legends

**Supplementary Table 1.** GenBank accessions of ITS and LSU sequences used for the phylogenetic analysis in this study. ITS and LSU rDNA sequences used in the phylogenetic analysis are listed with GenBank accessions and references. The strain used in this study are shown in bold.

| Species | Culture number | ITS | LSU | References |
| --- | --- | --- | --- | --- |
| *Teratoramularia infinita* | CBS 120815 | KX287544 | KX287248 | Videira et al. 2016 |
|  | CBS 141104; CPC 19488 | KX287545 | KX287249 | Videira et al. 2016 |
| *Teratoramularia kirschneriana* | CBS 113093; RoKi 1144 | GU214669 | GQ852627 | Crous et al. 2009 |
| *Teratoramularia persicariae* | CPC 11408 | KX287546 | KX287250 | Videira et al. 2016 |
|  | CPC 11409 | KX287547 | KX287251 | Videira et al. 2016 |
|  | CBS 141105; CPC 11410 | KX287548 | KX287252 | Videira et al. 2016 |
| *Teratoramularia rumicis* | NFCCI 5008 | MK967558 | MW276098 | Verma et al. 2021 |
| ***Teratoramularia rumicicola*** | **TR4** | **PX069134** | **PX069135** | **This study** |
|  | CPC 14652 | KX287549 | KX287254 | Videira et al. 2016 |
|  | CBS 141106; CPC 14653 | KX287550 | KX287255 | Videira et al. 2016 |
|  | CPC 14654 | KX287551 | KX287256 | Videira et al. 2016 |
| *Staninwardia suttonii* | CPC 13055 | DQ923535 | DQ923535 | Summerell et al. 2006 |

**Supplementary Figure 1.**
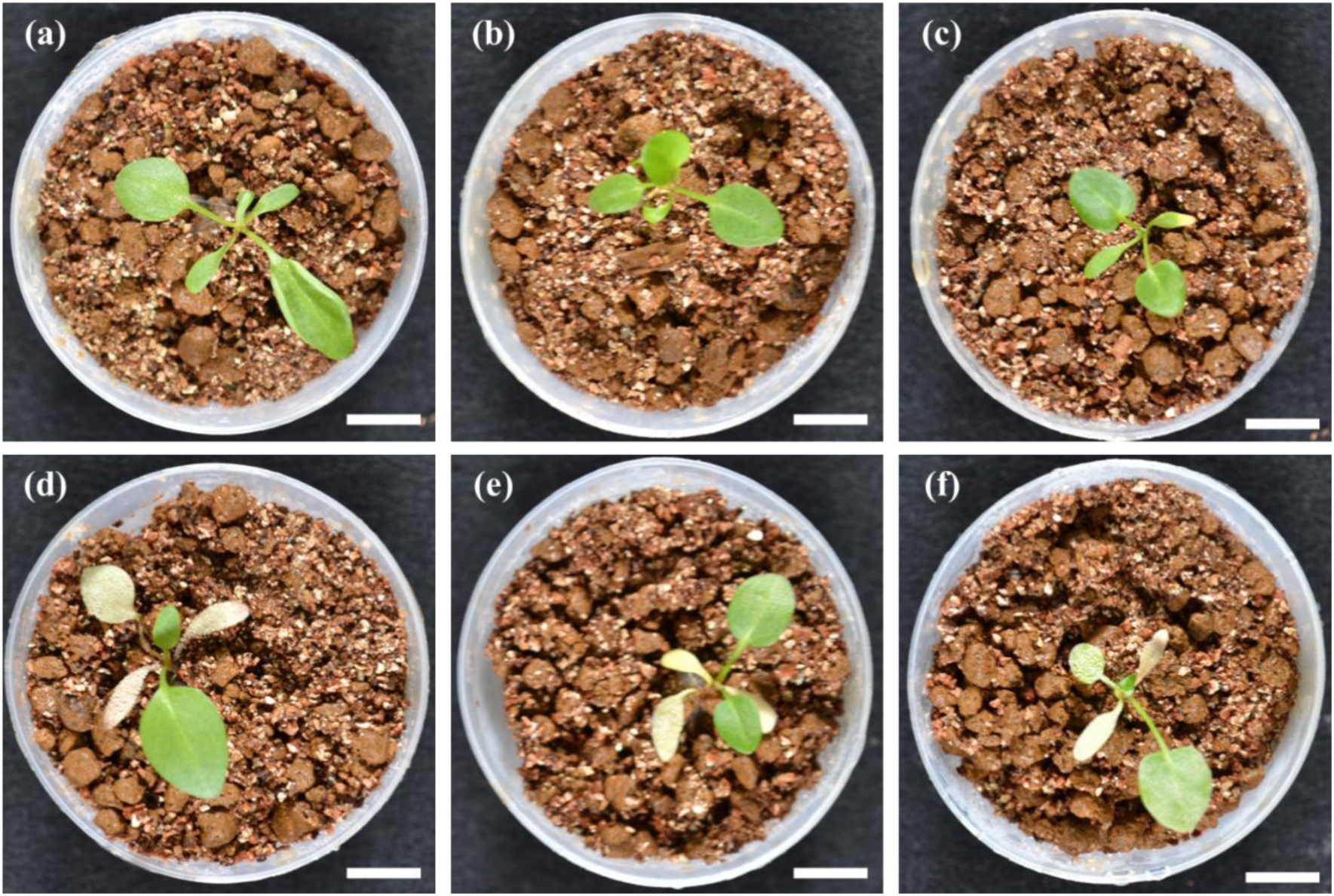
Pathogenicity of TR4 Against *Rumex* Species at 14 DAT. Foliar appearance 14 days after inoculation. (a–c) Untreated control plants of *R. crispus*, *R. japonicus*, and *R. obtusifolius*, respectively. (d–f) Corresponding TR4-inoculated plants showing visible white fungal growth and necrotic lesions. Scale bars = 10 mm

**Supplementary Figure 2.**
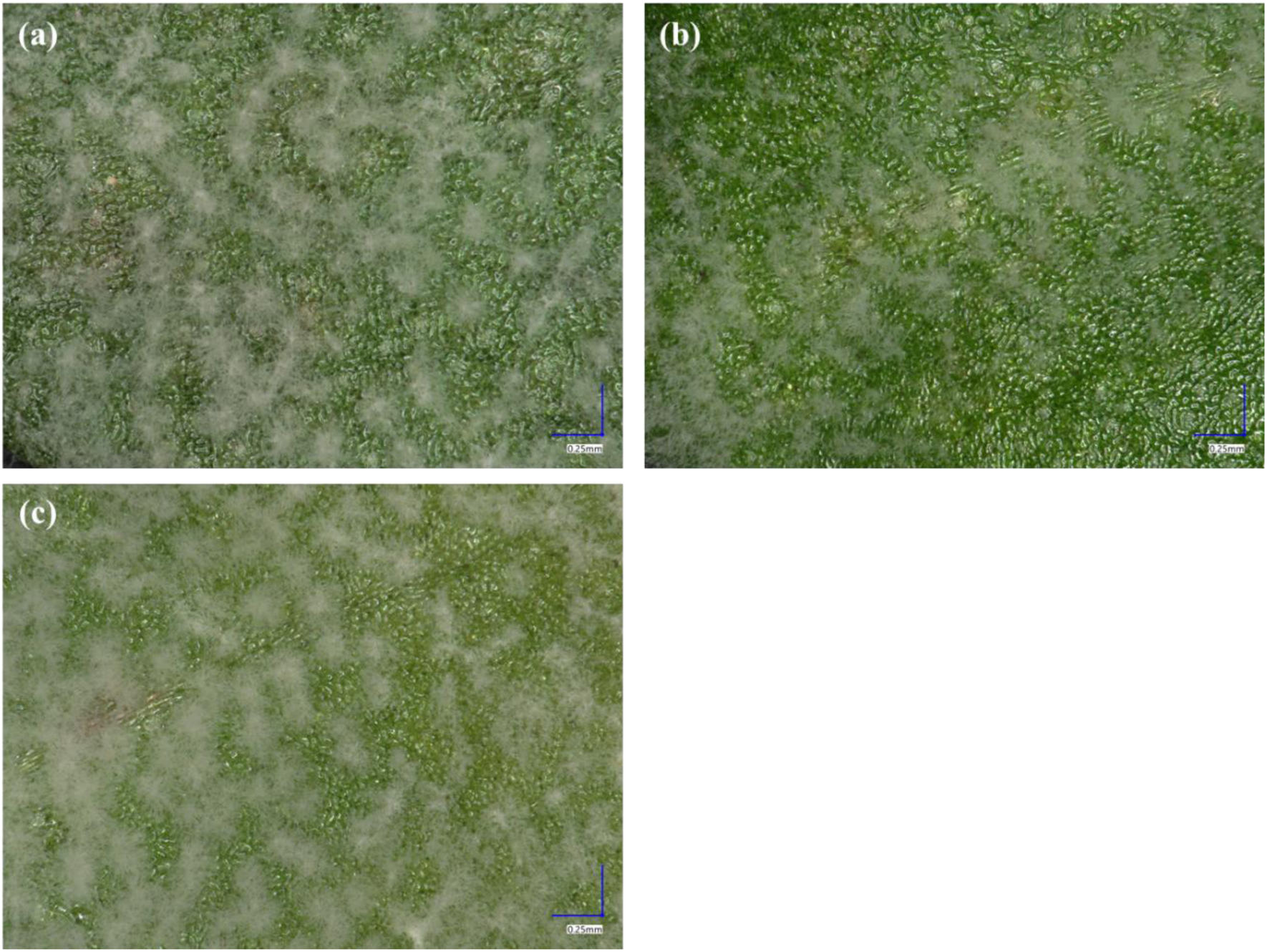
Digital microscope images showing white fungal growth of TR4 on the surface of infected *Rumex* leaves. Images represent the typical symptoms observed 21 days after inoculation. (a) *R. crispus*, (b) *R. japonicus*, (c) *R. obtusifolius*. Scale bars = 0.25 mm

## Notes

### Competing Interest Statement

The authors have declared no competing interest.

### Summary of Updates

Corrected italic formatting of scientific names in the title and abstract for accurate HTML display. No changes to PDF content.

